# Inferring super-resolved spatial metabolomics from microscopy

**DOI:** 10.1101/2024.08.29.610242

**Authors:** Luca Rappez, Kristina Haase

## Abstract

Current spatial metabolomics techniques have transformed our understanding of cellular metabolism, yet accessible methods are limited in spatial resolution due to sensitivity constraints. MetaLens, a deep generative approach, disrupts this trade-off by quantitatively propagating cellular-resolution *in situ* imaging mass spectrometry readouts to subcellular scales through integration with high-resolution light microscopy. MetaLens identifies subcellular metabolic domains with distinct molecular composition, enabling accessible label-free subcellular metabolomic analysis from microscopy.

## Main

Studies of metabolites and lipids in single organelles, including mitochondria^1^, endoplasmic reticulum (ER)^2^, and lipid droplets (LD)^3^, have broadened our knowledge of disease mechanisms^4,5^. While advancements in spatially resolved omics now allow detailed studies of RNA^6^ and protein distribution^7^, untargeted metabolomics *in situ* at similar spatial scales remains challenging. Currently, *in situ* metabolomics is prominently based on imaging mass spectrometry (IMS)^8^. In particular, matrix-assisted laser desorption/ionization (MALDI)-IMS enables soft ionization of endogenous metabolites, lipids, peptides, and proteins, revealing unprecedented single-cell molecular phenotypes ^8–11^. However, MALDI-IMS faces a trade-off between spatial resolution and sensitivity, limiting its application for high-throughput, wide-spectrum, and high-dynamic-range analysis^12^. This leaves an unmet need for accessible MALDI-IMS methods with subcellular resolution.

Deep learning has successfully been used to predict super-resolved omics readouts^13,14^, challenging traditional trade-offs between sensitivity and spatial resolution^15^. Inspired by these advances, we present MetaLens, a deep generative model that predicts spatially super-resolved MALDI-IMS metabolomic profiles from microscopy (**Fig. 1a**). Using a weakly supervised approach, MetaLens learns to propagate high-dynamic-range MALDI measurements to super-resolved pixels from microscopy data centered on sampled areas, marked by laser ablation marks (AMs). This enables MetaLens to locally estimate metabolite distributions within AMs at input microscopy resolution while predicting MALDI measurements. Post-training, MetaLens produces globally super-resolved images by virtually sampling microscopy pixels in sampled and unsampled areas, revealing subcellular structures unattainable at input MALDI resolution (**Fig. 1b**).

**Figure 1.**
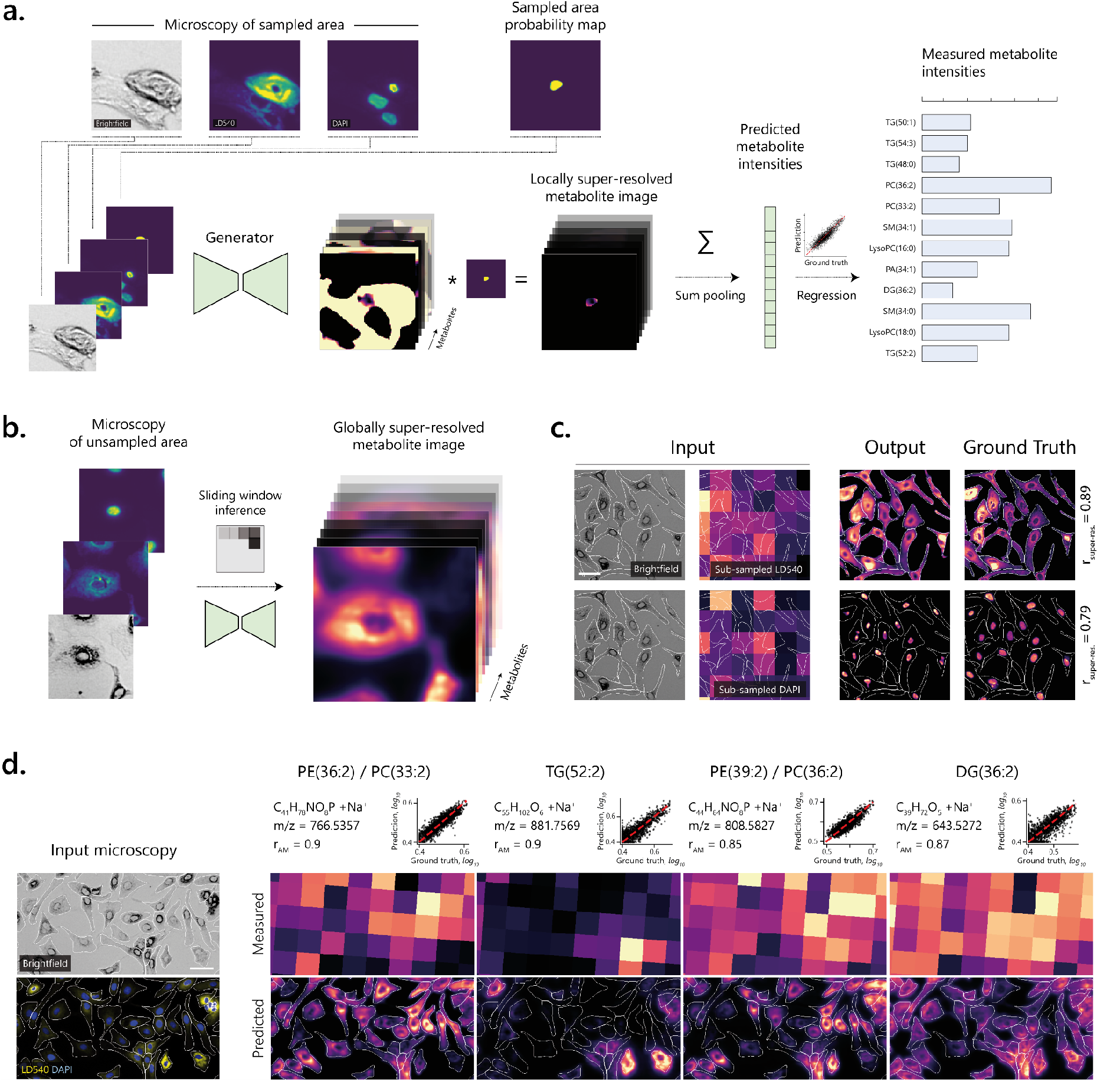
The MetaLens workflow, validation, and application to human hepatocytes. a, Deep generative model predicting metabolomics from microscopy while generating locally super-resolved metabolite images in sampled areas. **b**, After training, MetaLens produces globally super-resolved metabolite images of sampled and unsampled areas. **c**, Validation using synthetic pseudo-MALDI fluorescent data from LD540 (LDs) and DAPI (nuclei) staining. Brightfield images and pseudo-MALDI (input) predict super-resolved distributions (output) at native resolution (ground truth). r_super-res._ indicates pixel-level Pearson correlation between output and ground truth. Scale bar, 50 μm. **d**, Application to HepaRG hepatocytes with heterogeneous LD accumulation. Four representative lipids are shown (columns)(top row: sum formula, m/z, r_AM_: AM-level Pearson correlation, AM regression plot; red dashed line indicates perfect prediction), measured ion images (middle row), and predicted super-resolved ion images (bottom row). Scale bar, 50 μm.

We quantitatively evaluated MetaLens’ ability to super-resolve images from subsampled data. A synthetic pseudo-MALDI dataset was created from fluorescence images of nuclei (DAPI) and the neutral lipid dye LD540 (**Fig. 1c, Methods**). Mimicking sparse MALDI sampling, fluorescence intensities at native-resolution (0.7μm pixels) were summed within simulated 50μm-spaced AMs, creating low-resolution pseudo-ion images (50μm pixels). MetaLens was trained to predict these summed intensities from the brightfield channel and AM probability maps, then generate globally super-resolved images at microscopy resolution. MetaLens accurately predicted simulated AM intensities (DAPI: r_AM_=0.94, LD540: r_AM_=0.96) and generated super-resolved images closely matching original fluorescence distributions (DAPI: r_super-res._=0.79, LD540: r_super-res._=0.89). These results confirm MetaLens’ ability to reconstruct subcellular structures from input with over 70-fold lower spatial resolution.

Next, we applied MetaLens to a non-alcoholic fatty liver disease (NAFLD) model where human hepatocytes (HepaRG) are treated with oleic and palmitic acids to simulate steatosis, a condition characterized by excessive LD accumulation^16^. Using brightfield, LD540, and DAPI microscopy as input, we inferred super-resolution images of 72 lipids with a median r_AM_=0.79 (**Supplementary Fig. 1, Methods**). Several lipids showed particularly high prediction accuracies, including PE(36:2)/PC(33:2) (r_AM_=0.9), TG(52:2) (r_AM_=0.9), PE(39:2)/PC(36:2) (r_AM_=0.85), and DG(36:2) (r_AM_=0.87). Furthermore, MetaLens captured cell-cell variation, as shown by heterogeneous distributions of PE(36:2)/PC(33:2) and PE(39:2)/PC(36:2). Notably, TG(52:2) levels were elevated in LD540-positive cells, confirming the biological relevance of the readouts, as neutral lipids are expected to be LD-localized.

Next, MetaLens was challenged with generalizing metabolic quantitative predictions to an unseen test sample (**Supplementary Fig. 2**). Although the predictive accuracy was reduced relative to the validation set (unseen AM from training samples, r_AM-validation_=0.78, r_AM-test_=0.65), the predicted ion images closely matched the spatial patterns of the ground truth (median structural-similarity-index SSIM_test_=0.89, median peak-signal-to-noise-ratio PSNR_test_=31.15dB, **Methods**). MetaLens demonstrates robust generalization across biological replicates, capturing fundamental metabolic patterns across different samples of the same cell type under similar conditions.

Consistency of super-resolution inference was tested for 20 random AMs on a subset of cells and demonstrated that the run-to-run variability was highly conserved regardless of AM shape (median pixel-level Pearson correlation coefficient, r_PP_=0.94)(**Supplementary Fig. 3**). Inclusion of the AM brightfield channel reduced reproducibility across runs (median r_PP_=0.79), but maintained accuracy for individual AMs (median r_AM_=0.80). This suggests that brightfield does not provide additional context for estimating exact metabolite values, but reduces generalizability to unsampled areas, highlighting the importance of using the probability map to maintain global generalizability.

Principal Component Analysis (PCA) showed trends across the 72 predicted lipid distributions, at the super-resolved pixel level^17^ (**Fig. 2a, Methods**). Three prominent regions radiate from the nuclei, aligning with known^18^ lipid distributions: PC1 (red) correlates with cytosolic/perinuclear areas, PC2 and PC3 jointly cover near-nuclear areas, and PC3 extended cytoplasms. Distinct metabolic profiles were associated with different cellular compartments (**Fig. 2b**). The near-nuclear areas (green-turquoise, PC2/PC3) contains neutral lipids (DG, TG -consistent with perinuclear LD synthesis), sphingomyelins (SM -synthesized in the ER), and phospholipids (known membrane components of ER-Golgi). The perinuclear/cytoplasmic areas (PC1) are rich in phospholipids (PE, PC, PA), consistent with their known co-regulation and residence in ER, Golgi, and plasma membrane^18^.

**Figure 2.**
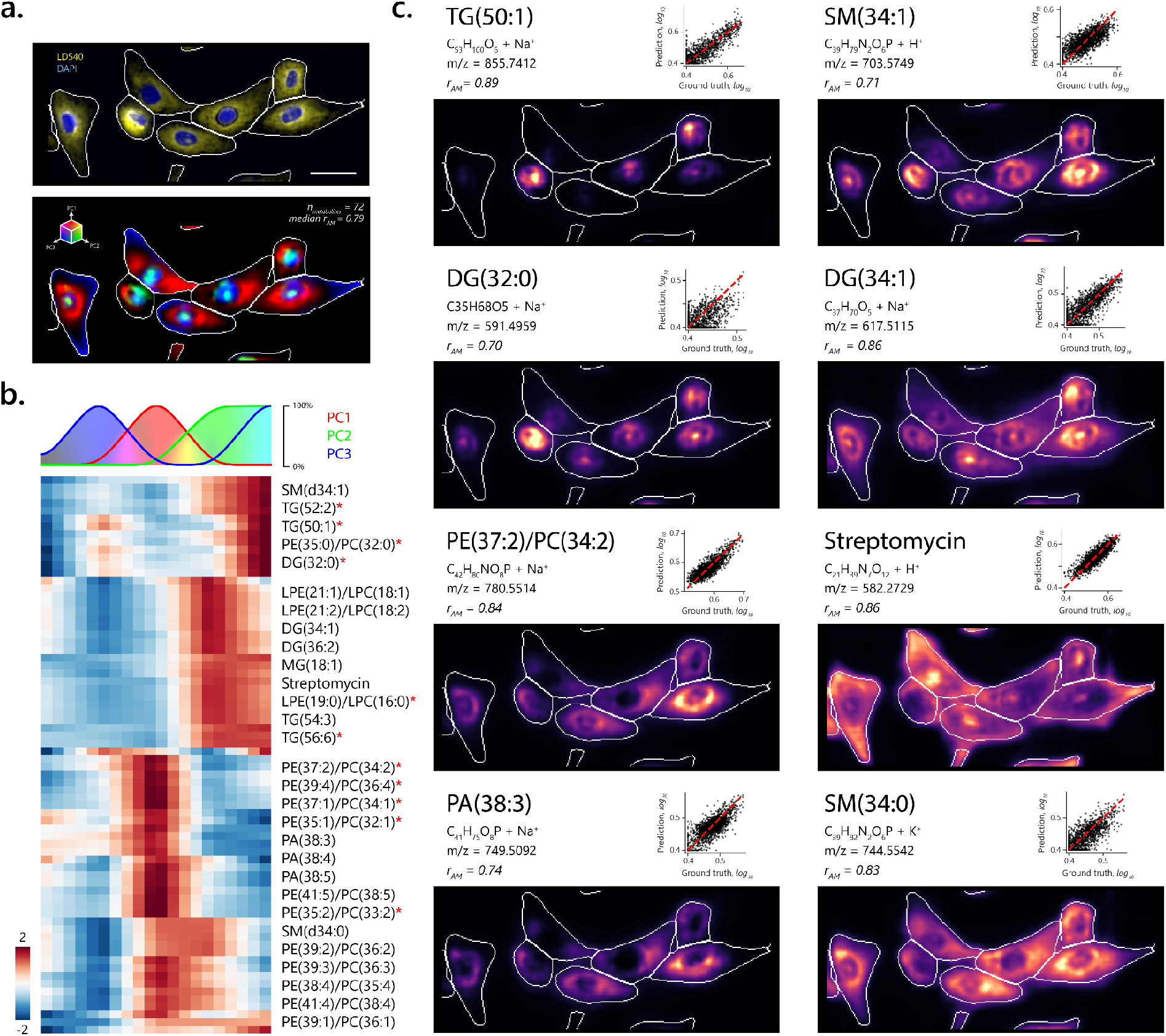
MetaLens reveals subcellular metabolic domains in human hepatocytes. a, Top: Microscopy image of dHepaRG hepatocytes stimulated with oleic and palmitic fatty acids (yellow, LD540; blue, Hoechst). Bottom: Spatial distribution of the three most prominent principal components (PCs) derived from 72 super-resolved metabolite images (n_metab._, predicted with median Pearson r_AM_=0.79). Scale bar, 50 μm. **b**, Decomposition of individual PC loadings identifies specific metabolic signatures corresponding to subcellular spatial domains. Color bar, metabolite-wise z-score. Red stars, metabolites previously validated by LC-MS/MS in mouse^9^. **c**, Representative subcellular distribution of metabolites, arranged from nuclear-localized to peripherally-distributed (top to bottom rows).

Notably, TG(50:1), a known marker of steatosis in HepaRG^9^, was perinuclear, indicating the likely presence of ER-bound or nucleoplasmic LD -a recently discovered phenomenon in certain cell types, including hepatocytes^19^ (**Fig. 2c**). Surprisingly, SM(d34:0) and SM(d34:1) display opposing cytoplasmic and nuclear/perinuclear localizations, suggesting a previously undescribed saturation-dependent compartmentalization^20^. Finally, streptomycin (from cell culture medium) shows a broad subcellular distribution, anti-correlating with LD540. This unexpected pattern could be due to mutual exclusion between LD540’s targets and streptomycin, which would have to be confirmed by further studies.

In summary, we have presented MetaLens, a powerful deep generative model for subcellular mass spectrometry that infers metabolome-wide super-resolved molecular distributions. Once trained on technical replicates, MetaLens predicts metabolic data from microscopy alone, enabling integration with other spatially resolved omics. By facilitating the quantitative analysis of multiple molecular readouts at subcellular resolution, MetaLens paves the way for an unprecedented understanding of health and disease across spatial scales.

## Online methods

### Image registration and segmentation

Pre- and post-MALDI microscopy images were co-registered using rigid body transformation (**Supplementary Fig. 4**). The acquisition area, encompassing all ablation marks (AMs), was manually cropped to yield a multi-channel image comprising both pre- and post-MALDI data. Cells were segmented using CellPose 2.0^21^ with brightfield images as input, resulting in the cell mask image Z. CellPose’s cyto2 model was fine-tuned on 20 randomly cropped fields of view with default hyperparameters. AMs were segmented using VascuMap’s method^22^, producing the AM probability map *P*.

### Microscopy-MALDI data integration

Inspired from ^9^, the AM mask *P*^*^was obtained from binarizing the AM probability map *P* ∈ [*0,1*] as: *P*^*^= *P* > *0*.*5* to measure the alignment angle *θ*^*^of the AM acquisition grid to the microscopy coordinate axes (**Supplementary Fig. 5a**). This angle was determined by minimizing the count of non-zero bins *b* in the orthogonal projections *O*:

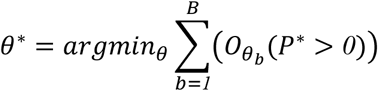

where *B* is the total number of bins and 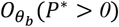 is the number of non-zero pixels from *P*^*^at the bin *b* following the orthogonal projection *O* at the angle *θ*.

This angle informed a Hough line transform to identify horizontal and vertical lines. These lines were filtered using K-means clustering, with cluster count matching the known number of rows or columns, yielding one line per column and row (**Supplementary Fig. 5b**). AMs were re-indexed by sorting columns and rows by their intercept parameters and examining intersections (**Supplementary Figs. 5c-d**).

To align AM coordinates with MALDI ion images, we determined the optimal homographic transformation (**Supplementary Fig. 6**). A sampling image *S* was created as:

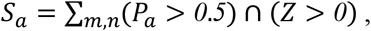

where *S*_*a*_ is the value for AM *a* of the sampling image *S* defined as the number of pixels (*m, n)* at the intersection of the mask *P*_*a*_ > *0*.*5* of the AM *a* and the cell mask *Z* > *0*.

Because the most abundant ions (originating from MALDI matrix salts) were found in extracellular spaces, the best transformation *T*^*^was found by minimizing the Spearman correlation *ρ* between the transformed sampling image *S* and the MALDI total ion count matrix *TIC* as: *T*^*^= *argmin*_*T*_*ρ*(*S*_*T*_, *TIC*), where *S*_*T*_ is the transformed sampling image and *TIC* is defined as: 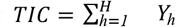, where *Y*_*h*_ is the MALDI ion image of ion *h* and *H* the total number of ions. This search considered all combinations of translations, flips, and 90-degree rotations.

### Training data generation

A multi-channel image I was constructed around each AM center:

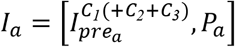

where 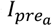 is the pre-MALDI microscopy, with channels *C*_*1*_(+*C*_*2*_ + *C*_*3*_), and *P*_*a*_ is the probability map of the AM *a*. Additionally, (+*C*_*2*_ + *C*_*3*_) indicates optional channels *C*_*2*_ and *C*_*3*_ as in the super-resolution validation experimentation where only the brightfield channel *C*_*1*_ was considered to predict fluorescence-derived pseudo-MALDI data (**Figure 1c**). To focus on the central AM *a*, the following steps were applied: binarization by thresholding the probability values at 0.5, region labeling by inspection of connected non-zero components, central region selection by computing the distance of each labeled component to the center of the image crop and dilation using a disk structural element of diameter 3. Further, AMs were filtered based on size to eliminate outliers and segmentation artifacts.

The MALDI ion image was normalized to the total ion count *TIC* and log-transformed channel-wise:

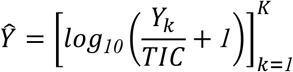

where 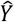 is the multi-channel normalized metabolite images, *Y*_*k*_ is the MALDI ion image for the metabolite *k* and *K* is the total number of predicted metabolites. The normalized metabolite images were further standardized channel wise as:

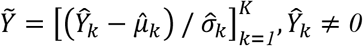

where 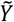 is the multi-channel standardized multichannel metabolite, 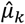 and 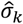 are the mean and standard deviations of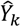, excluding zero values. The train-test split ratio was 90-10%.

### Model training

A DeepLabV3+^23^ model with a Resnet152^24^ encoder *G* was used as the image generator. The model transformation was defined as *G* : 𡄭^*M*×*N*×*C*^ → 𡄭^*M*×*N*×*K*^ where a multi-channel image of vertical and horizontal dimensions *M* and *N*, respectively, and *T* number of channels is transformed into a locally super-resolved metabolite image of matching spatial dimensions *M, N* and with *K* number of metabolites to predict. Image augmentation included: random affine transformations: scaling (0.75 to 1) and rotation (−45° to +45°), vertical and horizontal flips and transposition. The reflect border method was applied, requiring an additional central AM filter on the transformed probability map.

Inspired from^13^ and ^14^, the weakly supervised loss function *L* is expressed as:

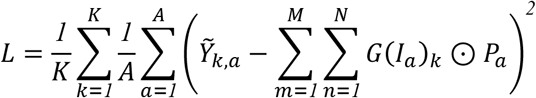

where *A* is the number of AMs in the whole image. *G*(*I*_*a*_)_*k*_ is the locally-super resolved metabolite image prediction for metabolite *k* by the generator model *G* from the microscopy image *I* centered on the AM *a*. 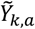 is the standardized measured abundance of the metabolite *k* at the AM *a*, ⊙ denotes element-wise multiplication.

Model performance was evaluated using Pearson correlation coefficient and determination coefficient r^2^ scores. Training used the AdamW optimizer with the following hyperparameters: batch size: 32, epochs: 200, initial learning rate: 1e-3, halved every 50 epochs. The model with the highest average Pearson correlation on the validation dataset was retained for inference.

#### Model inference

A sliding inference process iteratively generates image crop *I*_*w*_, at a window *w*, from a microscopy dataset 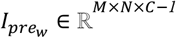, each appended with an arbitrary AM probability map *P*_*e*_ ∈ 𡄭^*M*×*N*×*1*^, giving: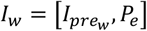, where *I*_*w*_ ∈ 𡄭^*M*×*N*×*C*^. The trained image generator *G* transforms each *I*_*w*_ into a multichannel image masked by the binarized AM probability map: *U*_*w*_ = *G*(*I*_*w*_) ⊙ (*P*_*p*_ > *0*.*5*), where *U*_*w*_ ∈ 𡄭^*M*×*N*×*K*^. A globally super-resolved image *U*_*global*_ is produced by averaging *U*_*w*_ over all windows 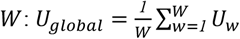. Finally, *U*_*global*_ is rescaled to natural normalized values producing 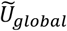as:

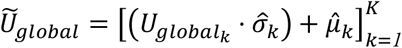

where 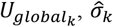 and 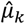 are the globally super-resolved image, the standard deviation and mean of the metabolite *k*, respectively. Both 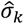 and 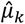 values are derived from the measured metabolite intensities used for training.

### Metabolite annotation and selection

Metabolite annotation was performed using the METASPACE^25^ cloud software (https://metaspace2020.eu) with the m/z tolerance of 3 ppm and false discovery rates of 50% against the Human Metabolome Database’s metabolite database v.2.5. All metabolites with a METASPACE FDR score inferior to 50%, detected for one of the samples used for training (link) and with a log_10_ median non-normalized abundance superior to 0.4 were retained for prediction, resulting in a group of 72 metabolites. A complete detail of the metabolite identification and validation of the metabolites listed in this study can be found in Supplementary Table 3 and Supplementary Data 1 of ref.^9^.

### Construction of the validation pseudo-MALDI dataset

To validate MetaLens, a subsampled pseudo-MALDI dataset mimicking the MALDI sampling process using LD540 and DAPI fluorescence channels was created. Subsampled data was generated by overlaying an arbitrary AM grid pattern on these fluorescence channels. For each AM, fluorescence intensities of pixels within the mark were summed and compiled into pseudo-ion images. Next, an image crop of the brightfield channel concatenated with the probability map of the arbitrary AM was generated. MetaLens was then trained to predict the summed LD540 and DAPI intensities for each AM. Using the same training and inference methods as for real MALDI data, MetaLens generated globally super-resolved images of LD540 and DAPI distributions. These reconstructed images were then compared to the original, full-resolution fluorescence channels allowing for quantitative, pixel-to-pixel comparisons between the MetaLens-generated super-resolved images and the ground truth high-resolution fluorescence data.

### Subcellular domain identification

To identify subcellular metabolic domains while minimizing the impact of cell-to-cell heterogeneity, we performed principal component analysis (PCA) at the super-resolved pixel level over the 72 predicted metabolite intensities for each cell independently. This approach allowed us to focus exclusively on intracellular subcellular domains. We visualized the spatial distribution of the first three principal components (PCs) within each cell using an RGB image: Red for PC1, Green for PC2, and Blue for PC3, with contrast adjustment for optimal visualization.

Given the heterogeneous lipid accumulation in the fatty-acid treated hepatocyte population, we observed variations in the spatial distribution and definition of PCs across cells. For simplicity and to establish a common reference frame, we manually selected a group of 35 stereotypical cells based on their similar spatial patterns (PC1 perinuclear, PC2 nuclear, and PC3 cytoplasmic). We then constructed a common PCA from the joint pixels of these 35 cells, rescaling the PC intensities cell-wise. This approach ensured consistent PC axes across those cells while highlighting the relative intensity of each metabolic domain within individual cells.

**Figure 2a-b** illustrates this method, showing the spatial distribution of PCs across cells and the relative correlation of each metabolite to the PCs based on their loading values. This approach allows for a robust comparison of subcellular metabolic domains across the heterogeneous cell population while preserving cell-specific variations and focusing on intracellular patterns.

## Data availability

All data used in this study are from the publicly available dataset of the peer-reviewed study by ^9^. This includes MALDI-imaging MS data, metabolite and lipid annotations, and microscopy datasets, which are accessible through METASPACE (https://metaspace2020.eu/project/Rappez_2021_SpaceM). The MALDI-imaging MS data are also available in the MetaboLights repository under accession number MTBLS78. Corresponding microscopy data can be found at the European Bioinformatics Institute BioStudies repository under accession number S-BSST369.

## Code availability

The code used to train the model as well as performing super-resolution inference will shortly be available at https://github.com/LRpz/MetaLens.

## Acknowledgements

We thank C. B. Vibe for editing the manuscript. We thank C. B. Vibe, T. Alexandrov, M. Stadler, W. Mirza, K. Stapornwongkul, M. Bernardello, P. Padmanaban, A. Akinbote, J. Trojanowski, D. Zawada, S. Owens, M. Shahraz and M. O. Ekelof for insightful discussions. This work received funding from the European Research Council (ERC) under Horizon 2020 research and innovation programme (Grant agreement No. 101040977).

## Author Contributions

L.R. conceived and implemented the method. L.R and K.H. wrote the manuscript. K. H. supervised the work.

## Supplementary figures

**Supplementary Figure 1.**
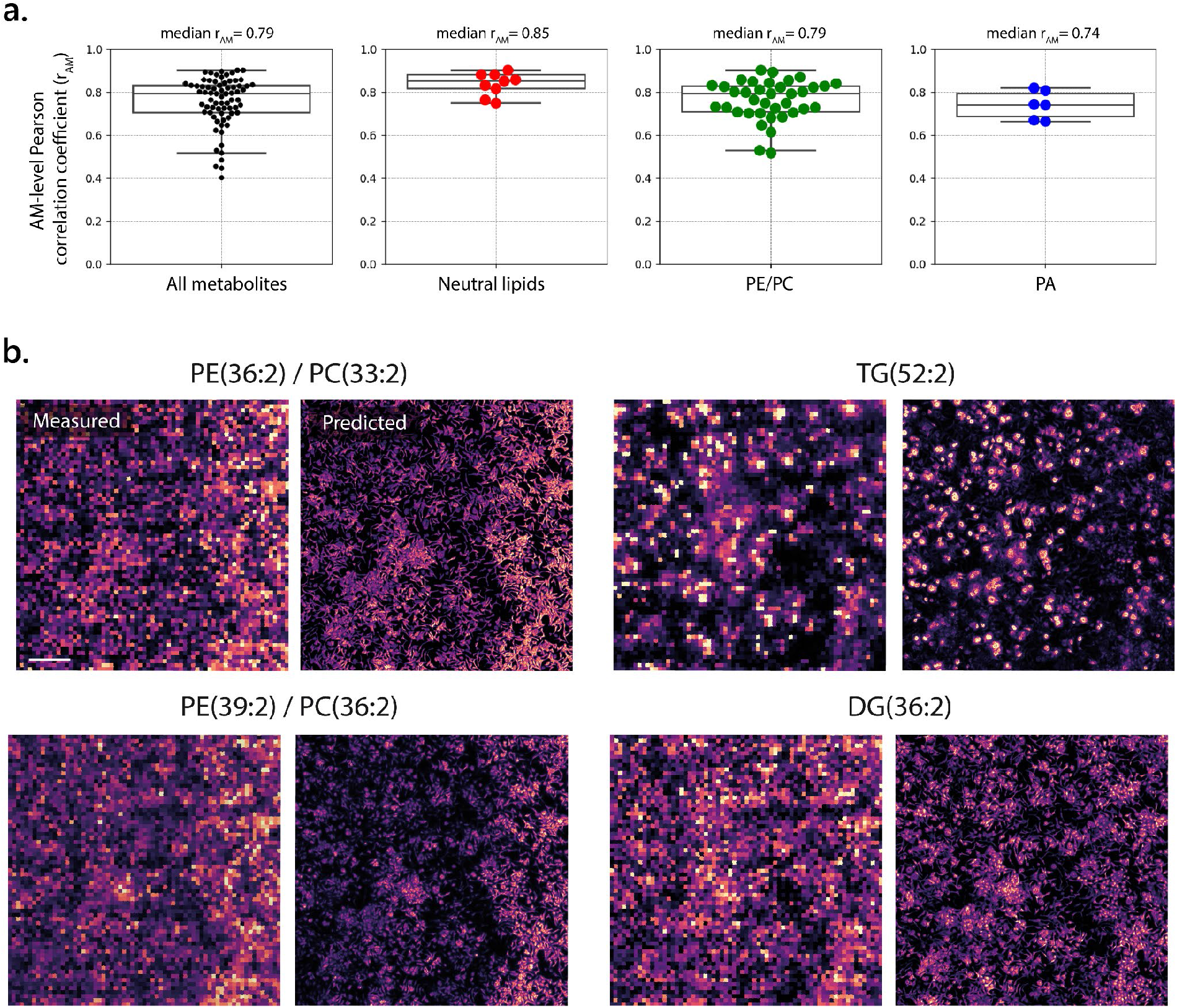
MetaLens prediction performance across metabolite classes. a. AM-level Pearson correlations (rAM) for 72 predicted metabolites. Black: all metabolites; red: neutral lipids (n=9); green: phosphatidylethanolamine/phosphatidylcholine (PE/PC) (n=37); blue: phosphatidic acid (PA) (n=6). Tukey box plots: center line, median; box limits, upper and lower quartiles; whiskers, 1.5× interquartile range. **b**. Comparison of measured and predicted super-resolved data across the sampled area. Scale bar, 500 μm.

**Supplementary Figure 2.**
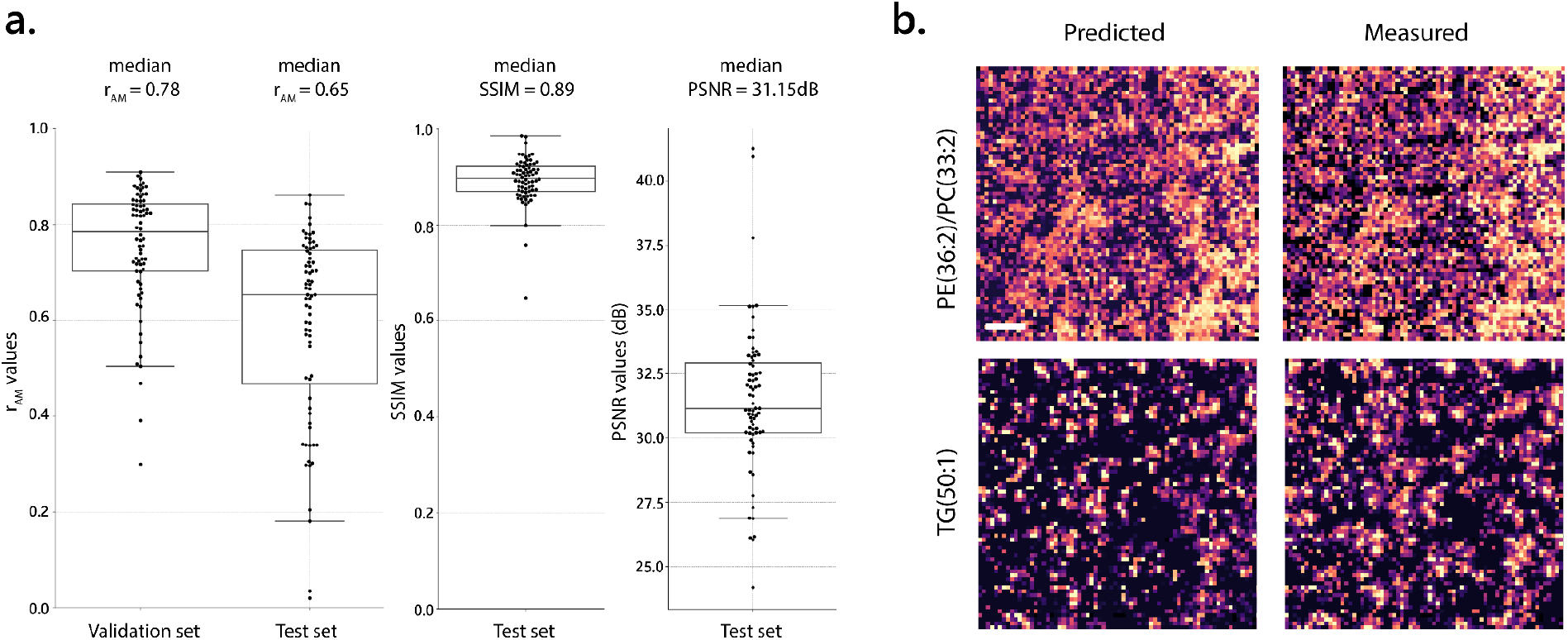
MetaLens performance on unseen data. a. Predictive accuracy metrics (r_AM_: AM-level Pearson correlation, SSIM: structural similarity index, PSNR: peak signal-to-noise ratio) for 72 metabolites in validation (unseen AMs, known samples) and test (unseen AMs, new sample) sets. Tukey box plots: center line, median; box limits, upper and lower quartiles; whiskers, 1.5× interquartile range. **b**. Predicted vs. measured ion-images (each pixel is an AM prediction) for the test sample. Scale bar, 500 μm.

**Supplementary Figure 3.**
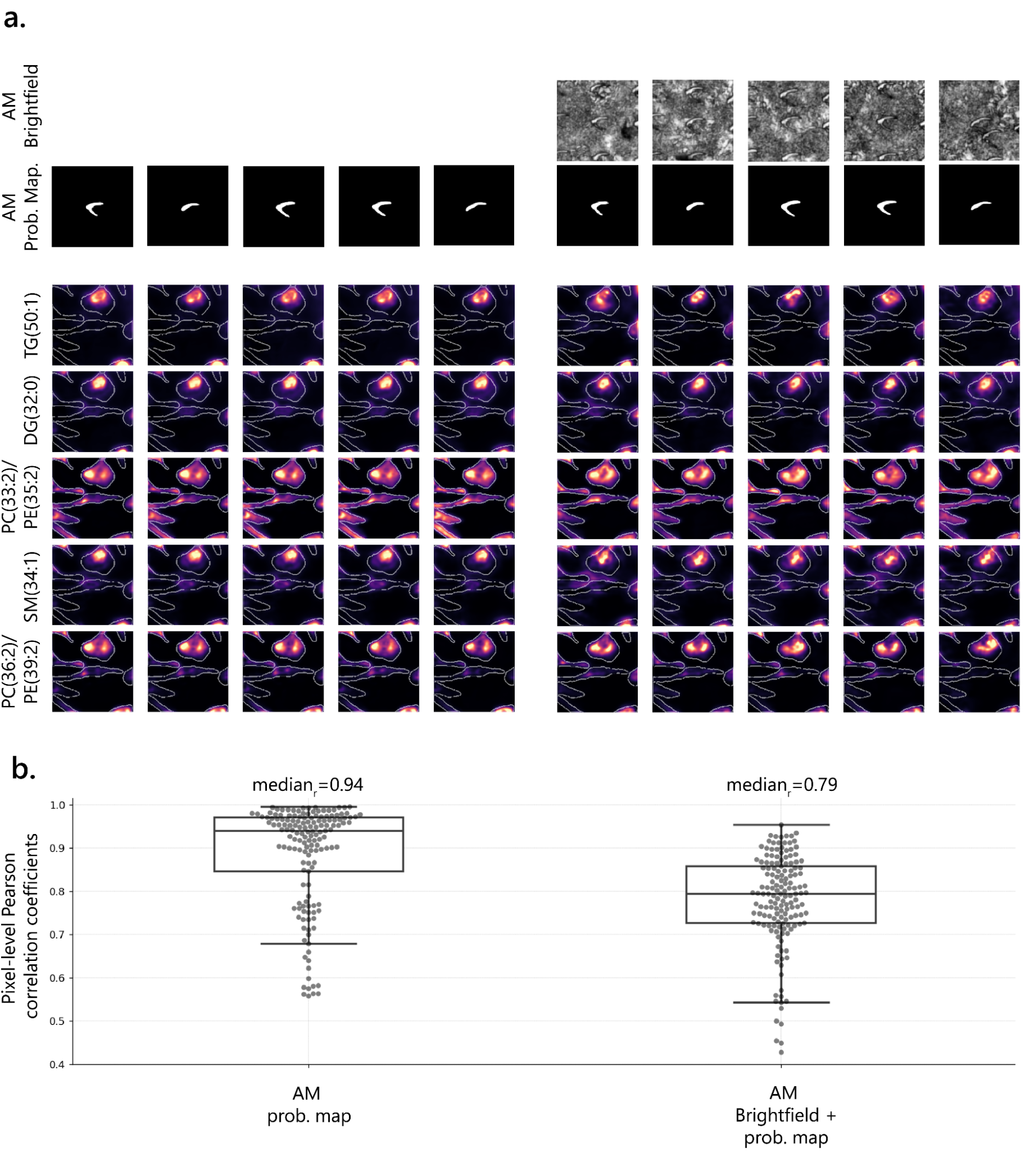
Effect of AM shape on inference variability. a. Subcellular lipid intensities (bottom 5 rows) predicted using different AMs (columns) during inference. Comparison between two models: one trained with only AM probability map (first 5 columns) and one with both brightfield and probability map as input (last 5 columns). **b**. Distribution of average pixel-level Pearson correlation coefficients between inferences from 20 randomly selected AMs for both models. Tukey box plots: center line, median; box limits, upper and lower quartiles; whiskers, 1.5× interquartile range.

**Supplementary Figure 4.**
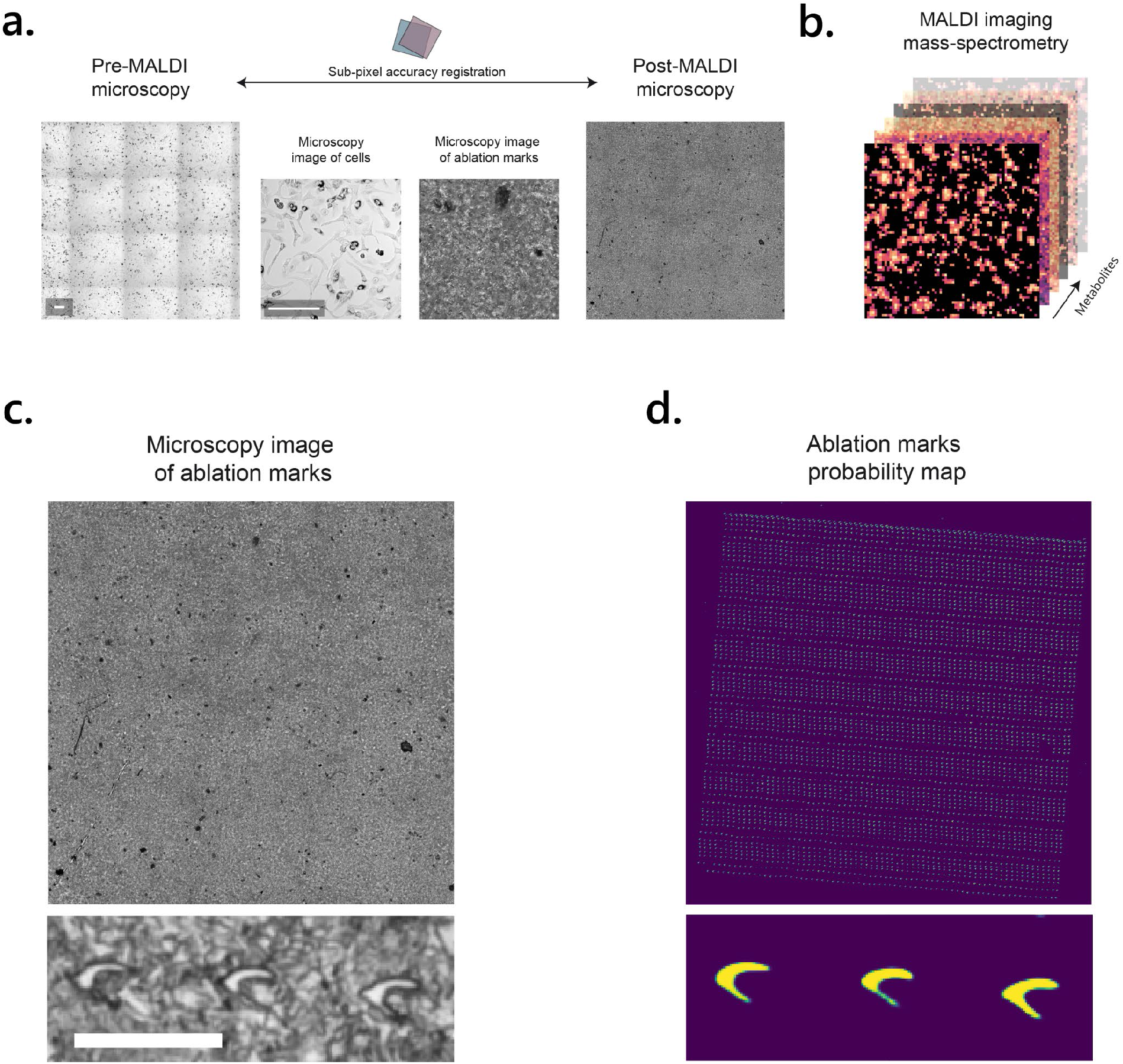
MALDI-microscopy integration and AM segmentation. a. Pre-MALDI microscopy image showing cells is registered with post-MALDI microscopy showing laser AMs. This precise alignment identifies sampled areas. Scale bar, 200 μm. **b**. Each laser shot produces a mass spectrum, which, when compiled over the acquisition grid, yields a multichannel ion image. **c-d**. AM segmentation using VascuMap’s neural network^22^, applied to brightfield images. The network’s final sigmoid layer output is a probability map (0-1 range) indicating likelihood of belonging to an AM. Scale bar, 50 μm.

**Supplementary Figure 5.**
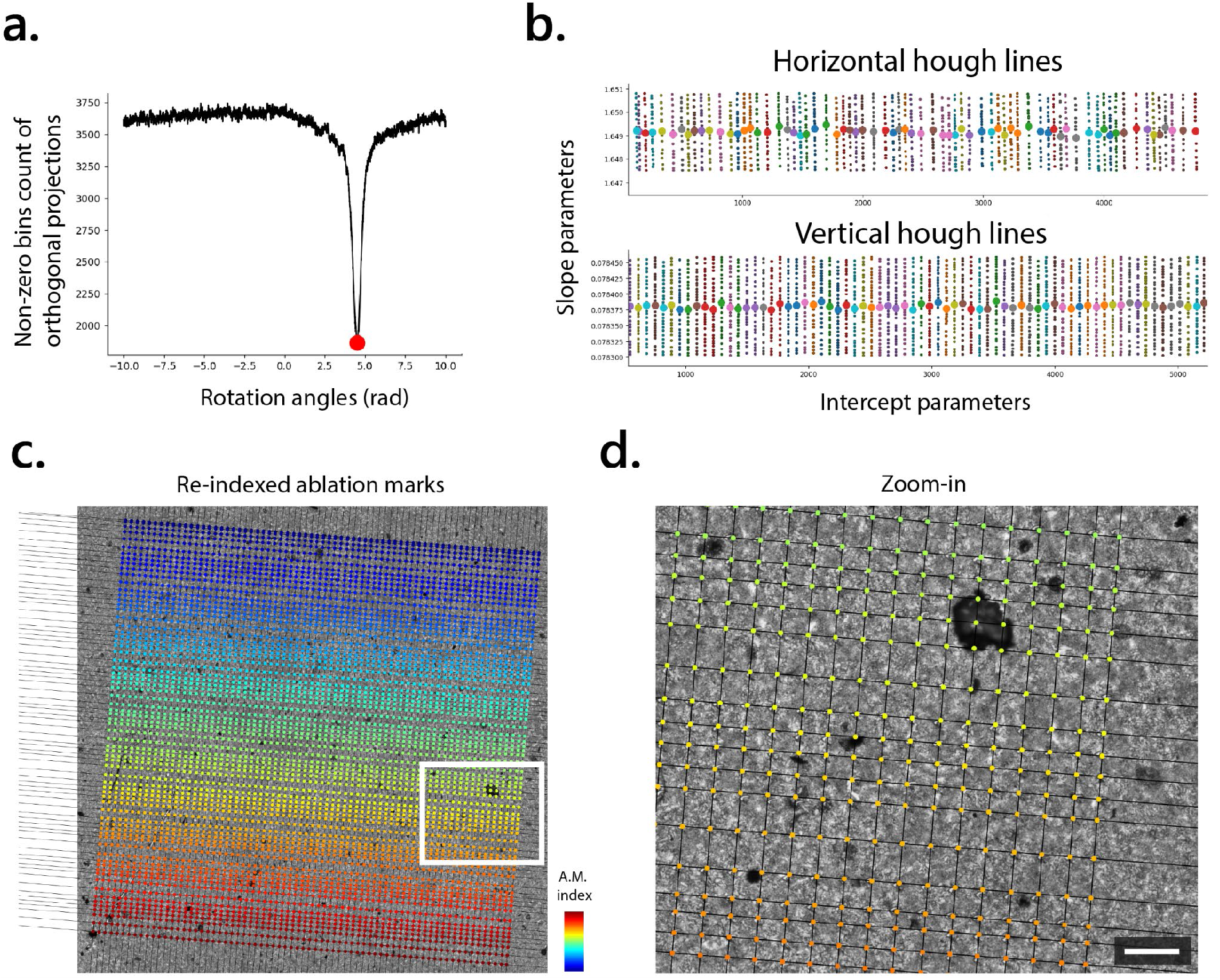
AM re-indexing. a. The AM grid acquisition angle is determined by minimizing non-zero bins in orthogonal projections of positive pixels from the probability map across various angles. The angle yielding the lowest non-zero bin count is considered the alignment angle relative to the microscopy coordinate system^9^.**b**. This angle informs a Hough line transformation to identify lines passing through the binarized AM probability map, initially producing multiple lines per row and column at +90 degrees. K-means clustering on alpha and intercept values isolates one line per row/column. **c-d**. AMs are reindexed by first sorting rows and columns by their intercept values, then iteratively examining intersections, mirroring the pixel arrangement in the ion image. Finally, each AM is assigned an index based on its closest intersection. Scale bar, 100 μm.

**Supplementary Figure 6.**
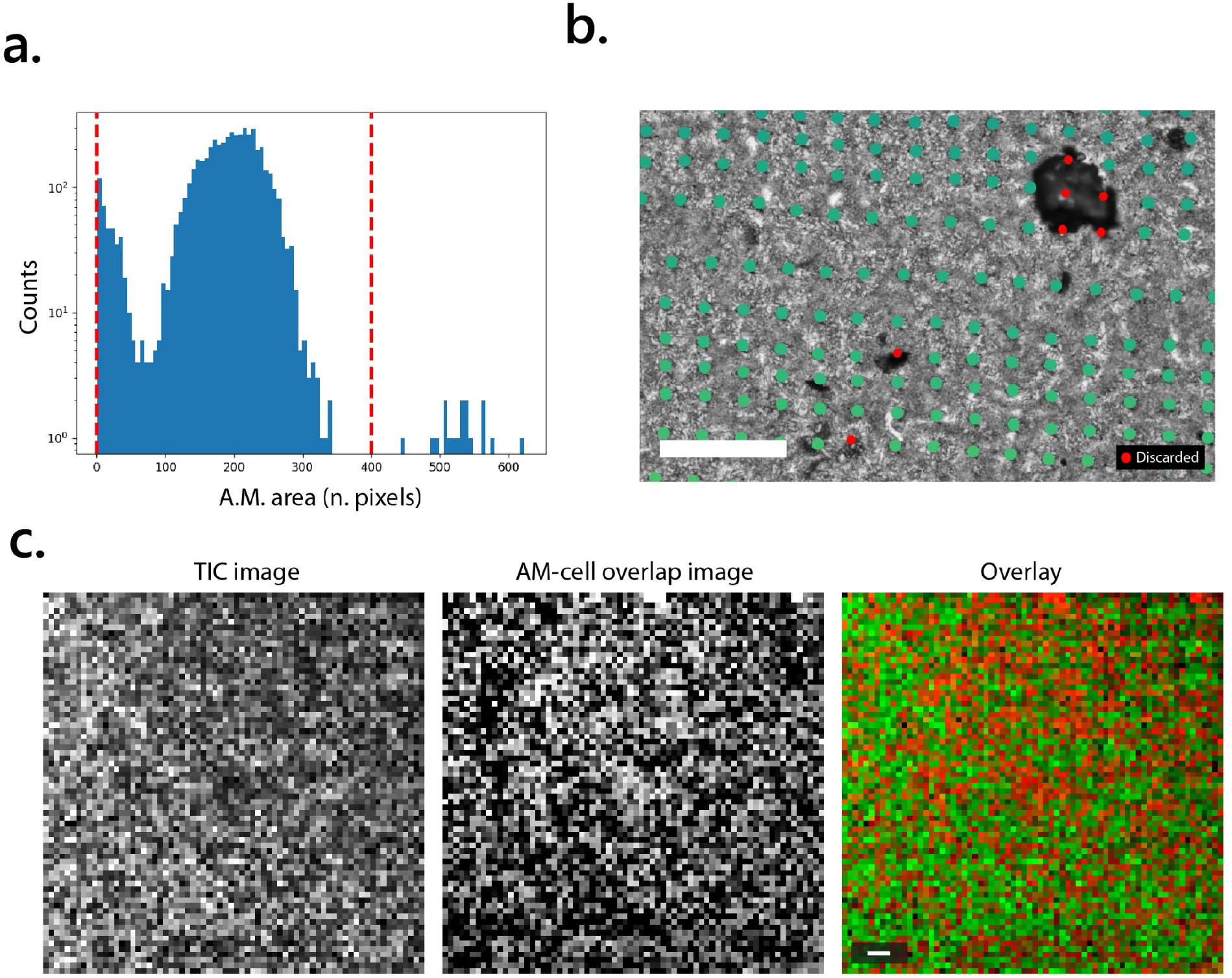
MALDI-microscopy integration refinement. a. Size filter applied to AMs to exclude segmentation artifacts and obstructed marks. **b**. Filter results; red dots indicate discarded AMs. Scale bar, 200 μm. **c**. Evaluation of ion image transformation to match microscopy. Total ion count (TIC) image is compared to AM-cell overlap image (pixel-wise overlap between AM and cell masks in microscopy space). High TIC is expected in cell-free areas due to the MALDI matrix, resulting in anticorrelation between TIC and AM-cell overlap images. The combination of flip, rotation, and transposition yielding the highest anticorrelation is deemed the correct transformation for integrating MALDI ion images with AM indices.

